# Dissociable pupil-linked arousal during overt and inner speech

**DOI:** 10.64898/2026.03.29.715074

**Authors:** Ryuta Kuwamizu, Ryoichi Watanabe, Nozomi Yamamoto, Kota Otani, Yusuke Moriguchi

**Author notes:** Correspondence should be addressed to: **Ryuta Kuwamizu, Ph.D.** Institute of Health and Sport Sciences, University of Tsukuba 1-1-1 Tennodai, Tsukuba, Ibaraki, Japan, **Ryoichi Watanabe, Ph.D.** Research Organization of Open Innovation and Collaboration, Ritsumeikan University, 2-150 Iwakura-cho, Ibaraki, Osaka 567-8570, Japan. Equal contributors (Ryuta Kuwamizu, Ryoichi Watanabe).

## Abstract

Inner (covert) speech, the silent production of language in one’s mind, plays a central role in human cognition and is known to activate speech-related cortical regions while also requiring executive and attentional resources. However, the conventional focus on cortical similarity or dissimilarity between inner and overt speech leaves unaddressed the question of whether arousal systems sustain cortical language processing. Here, using pupillometry across three controlled counting experiments, we show a dissociation between inner and overt speech. Overt speech consistently led to pupil dilation across all conditions. Inner speech, by contrast, produced a strikingly different profile: pupil constriction during simple counting, no deviation from baseline during noun-phrase counting, and clear dilation only under a high cognitive load, but with dilation remaining significantly less than that for overt speech in every condition. Covert verbal production thus proceeds at a reduced arousal cost, yet this cost scales flexibly with task demands rather than being fixed. This pupillometric dissociation reveals that inner and overt speech occupy fundamentally distinct arousal states, even when their cortical substrates partially overlap. These findings open a non-invasive window for tracking inner speech physiology across development, individual differences, and clinical populations in whom inner speech is altered.

## Introduction

There is an old saying that “the eyes speak as much as the mouth.” But do they speak the same language when we talk aloud versus when we talk silently to ourselves?

Inner speech, the act of speaking silently in one’s head, is a key cognitive tool for human thought and self-regulation^1^. Neuroimaging studies from the past two-three decades hold that inner speech, also referred to as covert speech, partially shares the same neural substrate as overt speech, particularly in left-lateralized language networks (e.g., Broca area)^1–3^. This has led to a prevailing motor-attenuation view positing that inner speech is essentially the same as overt speech, albeit with the final motor commands for articulation withheld^4^. However, this focus on cortical similarity or dissimilarity between inner and overt speech leaves a critical dimension unaddressed: the arousal systems that sustain cortical language processing. The lateral prefrontal cortex depends on monoaminergic and cholinergic neuromodulation for its proper function^5^, and these systems actively shape how cortical circuits process information—as shown in the auditory cortex, where arousal state determines the quality and reliability of sound-evoked neural responses^6,7^. A comparable dependence holds for language-related networks, where disruption of neuromodulatory input is associated with impaired language processing and poor recovery from aphasia^8^. Yet whether inner and overt speech differentially engage arousal systems has not been directly examined. We therefore hypothesized that arousal systems may fundamentally underlie any cortical similarity or dissimilarity between the two speech modalities.

Pupillometry provides a direct, non-invasive window into the arousal systems described above. Pupil diameter is a sensitive real-time index of arousal dynamics mainly linked to the locus coeruleus–noradrenergic and cholinergic systems ^9,10^, predicting aspects of subcortical and cortical arousal that heart rate or electrodermal measures do not directly reflect^9–15^. It is also subject to top-down executive control^16^ and reliably increases in response to cognitive effort, respiration, and physical activity^12,17–21^. Crucially, pupillometry requires no head-mounted devices and remains robust during overt articulatory movements, making it uniquely suited for directly comparing inner and overt speech within a single paradigm (in contrast to other brain imaging methods, which are often susceptible to significant speech-related noise^1,22^). Although pupillary responses during speech production have not yet been well studied, it has been suggested that overt vocalization elicits pupillary dilation^23,24^. Inner speech is known to recruit cortical processes that partly overlap with overt speech, including left-lateralized language regions^1,3^. Moreover, inner speech may place demands on sustained attention and verbal working memory, particularly when phonological representations must be internally maintained and monitored^1^. Some accounts further suggest that inner speech may also engage motor-inhibitory control to prevent overt articulation^4,25^. Additionally, motor imagery tasks have also been shown to elicit pupil dilation^26,27^, suggesting that pupil-linked arousal is engaged even when overt action is suppressed. On this basis, we hypothesized that pupil-linked arousal would also be engaged during inner speech, as a reflection of the cortical processing it requires.

In the present study, we addressed this question using a numerical counting paradigm. Counting provides a useful experimental model because it is a highly controlled verbal task with minimal affective variability, can be implemented in both inner and overt forms, and allows systematic manipulation of cognitive load while preserving the core structure of verbal production. It also makes it relatively feasible to experimentally verify, to some extent, whether inner speech is being engaged. Our recent work, for example, used numerical counting to show that preschool children (3-6 years old) are capable of inner speech to a measurable degree^28^. Although this paradigm does not capture the full range of naturally occurring inner speech, it reflects one form of inner speech that humans frequently use in everyday life^29^ and thus provides a useful model for investigating its physiological characteristics. We therefore first tested whether inner speech engages pupil-linked arousal in a manner comparable to overt speech, using a simple numerical counting paradigm (Experiment 1). Contrary to our primary hypothesis, inner speech produced not dilation but constriction relative to baseline, while overt speech led to dilation as expected, raising the question of whether this dissociation is a robust feature of covert verbal production or specific to the demands of simple numerical counting. We next examined whether the dissociation persists when verbal content is enriched: in Experiment 2, participants counted using grammatically structured numeral-noun phrases, a manipulation that increases phonological and morphological complexity while preserving the counting structure. Finally, in Experiment 3, we asked whether the dissociation holds under a higher cognitive load by requiring step counting, thereby examining whether inner speech arousal scales with task demands. This stepwise approach allowed us to characterize the conditions under which pupil-linked arousal during inner speech is attenuated relative to overt speech, and the extent to which that attenuation varies with verbal content and cognitive demands.

## Materials and Methods

### Participants

Across all three experiments, participants were healthy, native Japanese-speaking undergraduate and graduate students. The study was conducted in accordance with the Declaration of Helsinki and was approved by the Ethics Committee of the Unit for Advanced Studies of the Human Mind, Kyoto University. All participants provided written informed consent prior to participation.

In Experiment 1, 25 individuals were recruited, yielding a final sample of 23 (mean age = 21.65 years, SD = 3.02; 14 females, 9 males) after excluding 2 participants due to excessive missing data in the pupil recordings (see *Pupil Recording and Analysis*). In Experiment 2, 30 individuals were recruited, yielding a final sample of 27 (mean age = 20.23 years, SD = 2.05; 8 females, 19 males) after excluding 3 participants for the same reason. In Experiment 3, 30 individuals were recruited, and no exclusions were made, yielding a final sample of 30 (mean age = 21.97 years, SD = 6.37; 14 females, 16 males). The experimental design and analysis plan for Experiment 2 were preregistered on the Open Science Framework (OSF; https://osf.io/7efbv).

### Sample Size Determination

For Experiment 1, the sample size was determined using an a priori power analysis (G*Power) based on effect sizes reported in prior block-design pupillometry studies. We calculated that a minimum sample of 20 would be required to detect a large effect size (Cohen’s dz = 0.8)^19,30,31^ with 80% power at a Bonferroni-corrected α of 0.0167. To account for potential data attrition, we set our recruitment target at 25 participants. For the subsequent Experiments 2 and 3, we conducted a new power analysis based on the main effect size observed in our own Experiment 1 (Cohen’s d = 0.69, No-thought vs. Inner speech). This analysis indicated that a sample of 25 participants would be required to detect this effect with 80% power at the same Bonferroni-corrected α level. We therefore set our recruitment target at 30 participants for Experiments 2 and 3.

### Experimental Procedures

Across all experiments, participants completed a block-designed task while maintaining their gaze on an on-screen instruction panel. Each block consisted of a 30-s rest period followed by a 45-s task period (Figure 1). During the rest period, the word “Rest” was displayed on the screen. The first 15 s of the rest period served as a washout interval, and the last 15 s were used as the baseline period for pupillometry analyses. During both rest and task periods, participants were instructed to keep looking at the screen and to minimize head movements. During task periods, participants were presented with a list of three instructions (conditions) and were prompted with an arrow indicating which instruction to follow: (1) No-thought (attempt to think of nothing), (2) Overt speech (count aloud), and (3) Inner speech (count silently in the mind). One set comprised one block for each of the three conditions, and each participant completed three sets (i.e., three blocks per condition) (Figure 1A). The order of conditions was randomized. Pupil diameter was recorded continuously throughout. No explicit instruction was given regarding counting pace; participants counted at their own natural tempo. Compliance was assessed by asking participants to report the final number reached during the overt speech and inner speech conditions.

**Figure 1.**
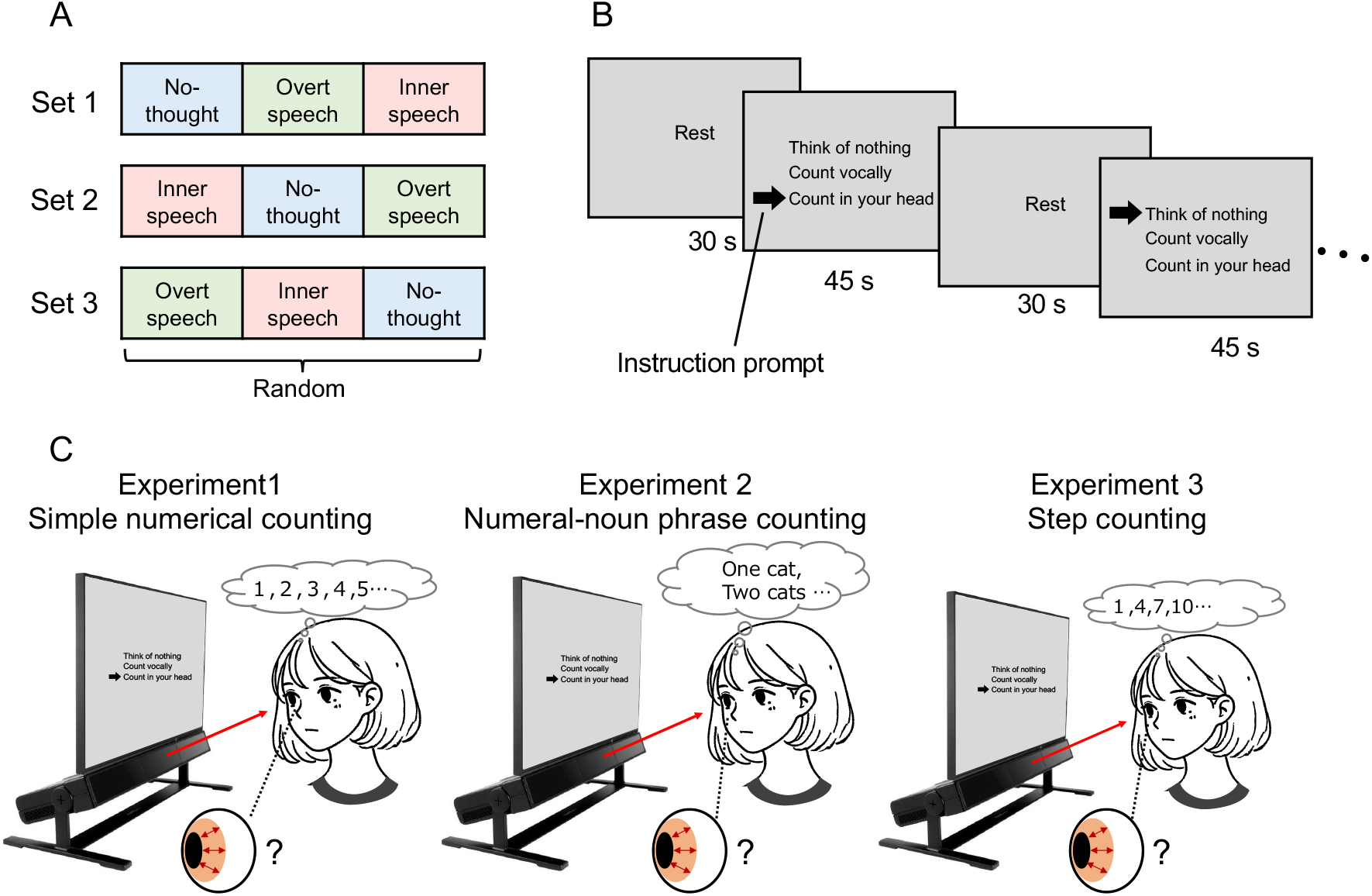
Schematic of the experimental paradigm and counting tasks. (A) Experimental session structure. Each participant completed three sets. Each set comprised one block of each of the three conditions (no-thought, overt speech, and inner speech). The order of the conditions was randomized within each set. (B) Trial sequence. Each block consisted of a 30-s rest period followed by a 45-s task period. During task periods, participants were prompted with an arrow indicating which instruction to follow: no-thought, overt speech, and inner speech. *(Note: Actual on-screen instructions were presented in Japanese: “休み” for Rest, “何も考えない” for No-thought, “声に出して数える” for Overt speech, and “心の中で数える” for Inner speech)*. (C) Experimental setup and task overview. Participants maintained their gaze on an on-screen instruction panel while pupil diameter was continuously recorded using a screen-based eye tracker. The illustrations depict the inner speech condition across the three experiments. In Experiment 1, participants counted upward from 1 using consecutive integers. In Experiment 2, participants counted in Japanese using grammatically structured numeral-classifier phrases (e.g., “one cat, two cats…”). In Experiment 3, participants performed step counting in increments of 2, 3, or 4 (an increment of 3 is illustrated here).

### Experiment 1: Simple numerical counting

Participants counted upward from 1 using consecutive integers (e.g., 1, 2, 3, 4…). Compliance questions were administered after each session.

### Experiment 2: Numeral-noun phrase counting

Participants counted in Japanese using grammatically structured numeral– classifier phrases that included the nominative particle *ga* (が) and category-specific classifiers. Before each session, the experimenter gave instructions about the semantic category (cats, balls, or books), and participants generated sequential counting phrases accordingly. For example, the animal category required phrases such as *neko ga ippiki, neko ga nihiki, neko ga sanbiki*… (“one cat, two cats, three cats…”), the object category required *booru ga ikko, booru ga niko*… (“one ball, two balls…”), and the book category required *hon ga issatsu, hon ga nisatsu*… (“one book, two books…”). This manipulation introduced not only semantic variation but also Japanese-specific morpho-phonological alternations across numbers (e.g., gemination and voicing in forms such as *ippiki*/*nihiki*/*sanbiki, ikko*/*niko, issatsu*/*nisatsu*), thereby increasing phonological/linguistic variability relative to simple numeral counting. The order of semantic categories across sessions was randomized and counterbalanced across participants. Compliance questions were administered after each block.

### Experiment 3: Step counting

Participants performed step counting. Before each session, the experimenter explained the counting rule: counting upward from 1 in increments of 2, 3, or 4 (e.g., 1, 3, 5, 7, 9…; 1, 4, 7, 10…; 1, 5, 9, 13…). The order of increment size across sessions was randomized and counterbalanced across participants. Compliance questions were administered after each block.

### Pupil recording and analysis

Pupil diameter was continuously recorded using a screen-based eye tracker (Tobii Pro Spectrum, Tobii AB, Danderyd, Sweden) at 60 Hz and processed in Tobii Pro Lab (Tobii Pro Spectrum, Tobii AB, Danderyd, Sweden). Visual stimuli were presented on a 23.8-inch (60 cm) monitor (FlexScan EV2451, EIZO, Japan). A 9-point calibration was performed at the beginning of each session. Participants were seated approximately 70 cm from the stimulus display and were instructed to maintain fixation and minimize head movements. Following previous works^19,20,30^, only valid samples from the right eye were analyzed; missing or invalid samples (e.g., due to blinks or gaze loss) were not included. To minimize luminance-related confounds on pupil size, visual parameters were strictly matched across the three conditions. All screens were presented on a uniform gray background (RGB: 120, 120, 120), and all task-relevant symbols/letters were rendered in black (RGB: 0, 0, 0). The on-screen instructions were presented in Japanese using the Meiryo UI font at size 48 (Figure 1B). The three conditions differed only in the on-screen position of the instruction arrow, ensuring that luminance and the amount of visual information on screen remained constant across conditions. For each block, a 30-s rest period preceded the task. The first 15 s of the rest period were treated as a washout interval, and the last 15 s were used to define the baseline. Baseline-adjusted pupil change (Δpupil diameter, mm) was computed as the mean pupil diameter during the subsequent 45-s task period minus the mean baseline value. Pupil data were downsampled to 1 Hz by taking the median within each 1-s bin, which reduces the influence of transient spikes and other brief artifacts on the summary time series. The 1-Hz time series was then averaged within each trial. Participants with insufficient valid pupil data were excluded (Experiment 1, n = 2; Experiment 2, n = 3; Experiment 3, n = 0): after 1-s binning, those with ≥50% missing samples across the session were removed from further analyses. Participants were also excluded if valid pupil data were entirely unavailable for any one condition. Illuminance in the testing room was monitored using a digital lux meter (Digital Lux Meter, model 78747; Shinwa Rules Co., Ltd., Japan). The light sensor was placed on the experimental desk with the receptor facing upward, and illuminance was maintained at 240–340 lux throughout the experiment.

### Analytic plan

To verify compliance with the inner-speech instructions, we computed Spearman’s *rho* between overt and inner speech final counts across participants for each experiment; a strong positive correlation would indicate that individual counting tempo was preserved across conditions. Separately, a paired *t*-test was conducted to assess whether the mean final count differed systematically between the two conditions.

For the primary pupillometry analysis, a one-way RM-ANOVA with condition (no-thought, inner speech, overt speech) as a within-subject factor and Δpupil diameter as the dependent variable was used to test whether conditions differed overall. When a significant main effect was observed, pairwise comparisons were conducted using Bonferroni-corrected paired *t*-tests across three planned contrasts: no-thought vs. inner speech, inner speech vs. overt speech, and no-thought vs. overt speech. One-sample *t*-tests against zero were additionally performed to determine whether each condition’s Δpupil diameter significantly deviated from the pre-task baseline (the 15 seconds before task onset). Significance threshold was *P* < 0.05 throughout. All statistical analyses were performed using GraphPad Prism 9. The analytic approach was established in Experiment 1, formally preregistered for Experiment 2 on the Open Science Framework prior to data collection (https://osf.io/7efbv), and applied without modification in Experiment 3.

## Results

### Validation of inner speech compliance

As an indirect check of compliance, we examined the correlation between the final numbers reported in the overt-and inner-speech conditions. Participants who counted relatively quickly during overt speech were expected to also do so during inner speech if they were engaging in the instructed covert counting process. Consistent with this, strong positive correlations were observed in Experiment 1 (*rho* = 0.92, *P* < 0.001), Experiment 2 (*rho* = 0.85, *P* < 0.001), and Experiment 3 (*rho* = 0.89, *P* < 0.001). Mean final counts (Inner vs. Overt speech) were 46.9 vs. 45.2 in Experiment 1 (*t*(22) = 0.70, *P* = 0.491), 33.6 vs. 29.7 in Experiment 2 (*t*(26) = 4.10, *P* < 0.001), and 100.4 vs. 101.7 in Experiment 3 (averaged across step sizes of 2, 3, and 4) (*t*(29) = 0.56, *P* = 0.578). Together, despite the absolute count difference in Experiment 2, these highly correlated findings provide indirect support for compliance with the inner-speech instructions.

### Pupillometry

### Experiment 1: Simple numerical counting

Baseline-adjusted pupil diameter differed significantly across conditions (repeated-measures one-way ANOVA: *F*(1.48, 32.65) = 8.92, *P* = 0.002) (Figures 2A and 2B). Contrary to our initial hypothesis, pupillary responses during inner speech were significantly smaller than those during no thought (*t*(22) = 3.32, adjusted *P* = 0.009) and overt speech (*t*(22) = 3.51, adjusted *P* = 0.006), whereas no significant difference was observed between no thought and overt speech (*t*(22) = 1.91, adjusted *P* = 0.208). To clarify the direction of these effects relative to baseline, one-sample tests showed that overt speech elicited significant pupil dilation (*t*(22) = 2.15, *P* = 0.043), whereas inner speech elicited significant pupil constriction (*t*(22) = -2.93, *P* = 0.008). The no-thought condition did not differ from baseline (*t*(22) = 0.70, *P* = 0.490).

**Figure 2.**
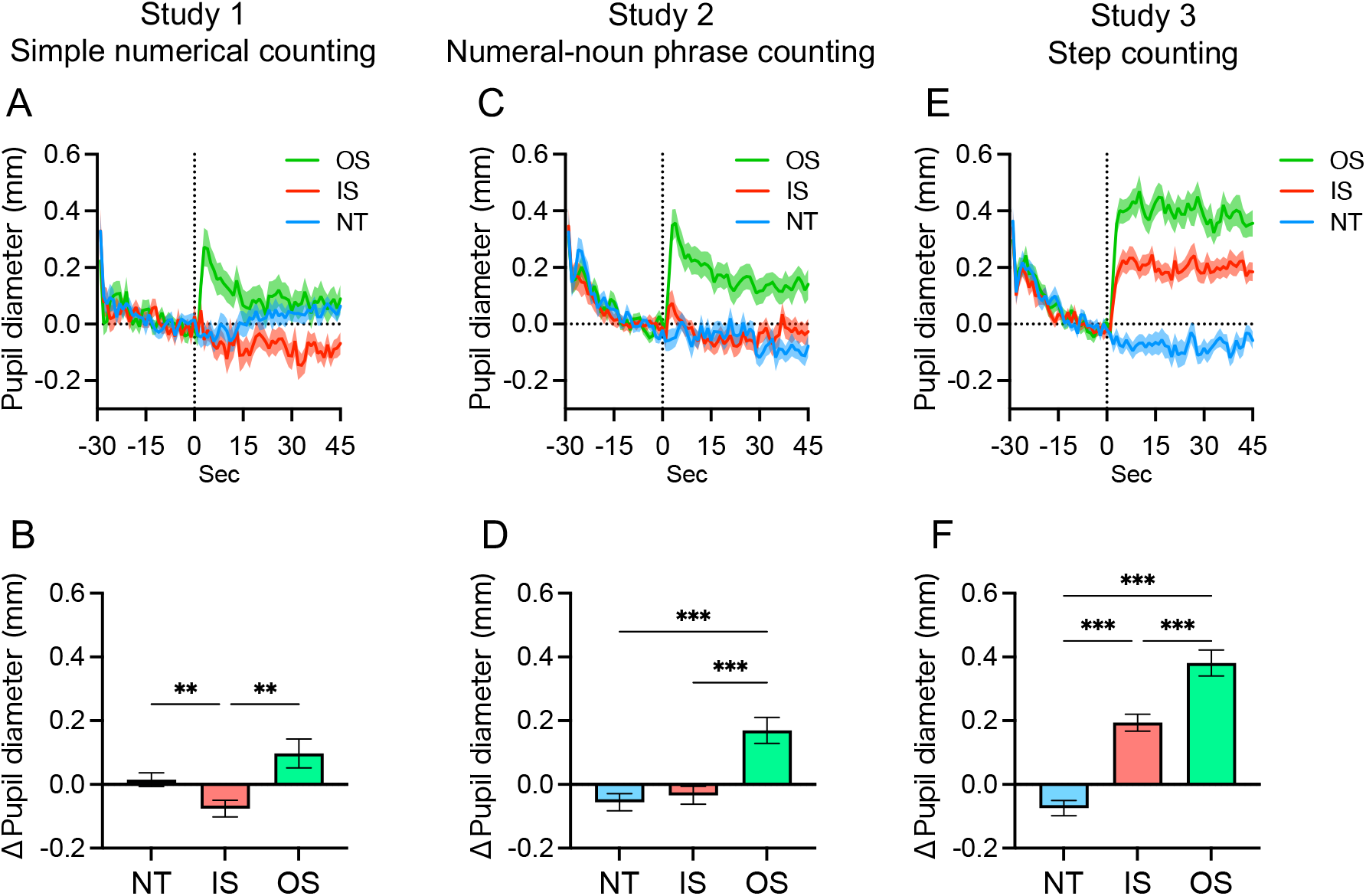
Pupil change between inner and overt speech across experiments. (A, C, E) Time courses of baseline-adjusted pupil diameter across the 30-s rest and 45-s task periods. The vertical dotted line at 0 s indicates the onset of the task. Solid lines represent the mean pupil diameter for the overt speech (OS, green), inner speech (IS, red), and no-thought (NT, blue) conditions. Shaded areas indicate the standard error of the mean (SEM). Data represent Experiment 1 (simple numerical counting) (A), Experiment 2 (numeral-noun phrase counting) (C), and Experiment 3 (step counting) (E). (B, D, F) Mean baseline (15 s pre-task)-adjusted pupil change (ΔPupil diameter) during the 45-s task period for Experiment 1 (B), Experiment 2 (D), and Experiment 3 (F). Error bars represent SEM. Asterisks indicate significant differences between conditions assessed via post hoc paired-samples *t*-tests following a significant repeated-measures ANOVA (** *P* < 0.01, *** *P* < 0.001).

### Experiment 2: Numeral-noun phrase counting

Baseline-adjusted pupil diameter differed significantly across conditions (repeated-measures one-way ANOVA: *F*(1.48, 38.53) = 18.91, *P* < 0.001) (Figures 2C and 2D). Consistent with Experiment 1, pupil size during overt speech was significantly larger than that during no thought (*t*(26) = 4.45, adjusted *P* < 0.001) and inner speech (*t*(26) = 5.41, adjusted *P* < 0.001). However, contrary to our pre-registered hypothesis (https://osf.io/7efbv) based on Experiment 1, inner speech and no thought did not differ significantly (*t*(26) = 0.71, adjusted *P* > 0.999). To clarify the direction of these effects relative to baseline, one-sample tests showed that overt speech elicited significant pupil dilation (*t*(26) = 4.13, *P* < 0.001), whereas no thought showed a small but significant pupil constriction (*t*(26) = -2.06, *P* = 0.049). Inner speech did not significantly differ from baseline (*t*(26) = -1.22, *P* = 0.234).

### Experiment 3: Step counting

Baseline-adjusted pupil diameter differed significantly across conditions (repeated-measures one-way ANOVA: *F*(1.53, 44.42) = 56.67, *P* < 0.001) (Figures 2E and 2F). During counting with step sizes of 2, 3, and 4, pupillary responses differed across all three conditions: inner speech responses were significantly larger than those during no thought (*t*(29) = 6.75, adjusted *P* < 0.001) but significantly smaller than those during overt speech (*t*(29) = 5.57, adjusted *P* < 0.001), and overt speech also elicited significantly larger responses than no thought (*t*(29) = 8.55, adjusted *P* < 0.001). To clarify the direction of these effects relative to baseline, one-sample tests showed that no thought elicited a small but significant pupil constriction (*t*(29) = -3.13, *P* = 0.004), whereas both inner speech (*t*(29) = 7.31, *P* < 0.001) and overt speech (*t*(29) = 9.33, *P* < 0.001) elicited significant pupil dilation.

## Discussion

The results of the present study do not support the initial assumption that inner and overt speech produce equivalent pupillary responses. Across three experiments, overt speech consistently elicited pupil dilation, whereas inner speech invariably produced reduced responses relative to overt speech. Critically, the direction of the inner-speech pupillary response relative to baseline was not fixed: pupil constriction occurred during simple numerical counting (Experiment 1), no significant deviation from baseline occurred during noun-phrase counting (Experiment 2), and clear dilation occurred during cognitively demanding step counting (Experiment 3). Together, these findings indicate that inner-speech-linked pupil responses are robustly attenuated relative to overt speech, while their direction varies systematically with both the specific verbal content and the cognitive demands of the inner-speech task.

A fundamental challenge in inner-speech research is verifying that participants are actually engaged in covert verbal production. The numerical counting paradigm offers an indirect but meaningful compliance check: participants who count rapidly during overt speech also tend to do so during inner speech. In comparing counting speed between inner- and overt-speech tasks, our task-compliance results were consistent with this tendency (*rho* > 0.85 across all three experiments).

The most robust finding across all three experiments was the consistent pupil dilation elicited by overt speech, suggesting that the presence of vocalization is a critical determinant of pupil-linked arousal. Pupil diameter is a well-established index of cognitive effort and brain arousal state^11,17^, reflecting coordinated activity across the locus coeruleus–noradrenergic system^15,32^, the basal forebrain cholinergic system^10^, the orexin system^33^, and associated cortical networks^13,14^. Although pupillometry has been applied to memory load^17^, lexical processing^34^, speech planning^34^, vocal reading^23,24^, musical imagery^35^, and physical activity^19^, direct comparisons of inner speech, overt speech, and instructed no-thought within a matched task structure are, to our knowledge, absent. The consistent overt-speech-induced pupil dilation observed here likely reflects the integrated arousal recruitment associated with motor execution, respiratory variation, sensorimotor feedback, and attention^15,18,24,36^, each of which is presumed to engage brainstem arousal systems and cortical networks^13,36^, potentially contributing to efficient speech production^5,8^. Inner speech, by contrast, did not approach overt-speech levels of pupil dilation in any condition, suggesting that covert verbal production can be executed without recruiting pupil-linked arousal systems and cortical networks to the same degree.

Interestingly, inner speech was not uniformly associated with low pupil-linked arousal. Under a minimal cognitive load (Experiment 1), inner speech produced pupil constriction even relative to the no-thought condition, suggesting that highly routinised covert counting is executed in a markedly low-arousal state. As cognitive demand increased (Experiment 2), this constriction was abolished, and inner speech no longer differed from baseline. Under the highest load (Experiment 3), inner speech produced clear dilation, converging toward—but remaining significantly below—that of overt speech. Inner speech therefore maintains a consistently attenuated response profile relative to overt speech while tracking cognitive load in a qualitatively similar manner.

Past neuroimaging studies have established that inner and overt speech share language-related networks and speech production processes^2,37^, and inner speech has recently been theorized as a form of inhibited overt speech^4^. Our findings do not challenge this shared neural substrate; rather, they reveal that, at the level of pupil-linked arousal, inner speech operates in a distinct mode that is dissociated from the obligatory arousal accompanying voice production. Inner speech may therefore utilize overlapping neurocognitive mechanisms while constituting verbal processing at reduced physiological cost. This interpretation aligns with the observation that inner speech can proceed faster than overt speech^38^ and with the Experiment 2 finding that participants counted to higher numbers during inner speech within the same time window, indicating more efficient progression.

Given that pupillometry is non-invasive and contactless, these findings open new avenues for tracing the Vygotskian internalizations of overt into inner speech across development and for examining individual differences in inner speech physiology in naturalistic settings. The capacity of inner speech to sustain verbal processing at reduced arousal cost may be advantageous where efficient cognitive activity and mental calm must coexist—as in cognitive control^39^, sport performance^39^ or routinised inner counting as a sleep aid^40^—yet may also enable maladaptive cognitive loops that contribute to rumination^41^. Conversely, the elevated arousal accompanying overt speech may carry functional value: Fawcett et al.^23^ showed that the pupil dilation elicited by reading aloud correlates with the memory advantage for spoken over silent reading words, suggesting that overt speech leverages its greater arousal cost to strengthen encoding. Together, these asymmetric profiles point toward translational applications of pupillometry at the intersection of speech modality and subcortical and cortical arousal regulation^39^.

Several limitations warrant consideration. First, the inner speech induced here was task-evoked and numerically constrained; it does not generalize to spontaneous or dialogic inner speech, which represent distinct and richer forms of covert verbal experience^42,43^. Second, the no-thought condition reflects an instructed attempt to suppress thought rather than verified mental silence; mind-wandering cannot be entirely eliminated, and this condition should be regarded as a tentative resting baseline^37^. Nevertheless, the replicability of the dissociation across three independent experiments, each employing a different counting paradigm, supports the robustness of the present findings as a foundational physiological account of inner versus overt speech.

In conclusion, inner and overt speech are characterized by dissociable pupil-linked arousal profiles. Overt speech consistently induced pupil dilation, whereas inner speech produced uniformly attenuated responses whose direction, constriction, no change, or dilation, depended on the verbal contents of the inner speech. This dissociation indicates that the pupil-linked arousal states underlying covert verbal production are fundamentally distinct from those engaged during overt voice production.

## Credit author statement

**Ryuta Kuwamizu:** Conceptualization, Methodology, Performing experiments (Experiment 1), Data analysis, Visualization, Writing – Original draft preparation, Project administration, Funding acquisition; **Ryoichi Watanabe:** Conceptualization, Methodology, Performing experiments (Experiments 2 & 3), Writing – Reviewing and editing, Funding acquisition; **Nozomi Yamamoto**: Writing – Reviewing and editing, Participant recruitment, Funding acquisition, **Kota Otani**: Conceptualization, Performing experiments (Experiment 1, assistance), Writing – Reviewing and editing, Funding acquisition; and **Yusuke Moriguchi**: Conceptualization, Supervision, Writing – Reviewing and editing, Funding acquisition.

## Data availability

The source data underlying the figures and analyses will be deposited on the Open Science Framework and made publicly available upon journal publication.

## Acknowledgements

The authors express their gratitude to Ms. Noguchi (ELCS English Language Consultation Services, Japan) for helping with the manuscript.

